# Sonlicromanol improves neuronal network dysfunction and transcriptome changes linked to m.3243A>G heteroplasmy in iPSC-derived neurons

**DOI:** 10.1101/2020.08.11.246140

**Authors:** T.M. Klein Gunnewiek, A. H. A. Verboven, M. Hogeweg, C. Schoenmaker, H. Renkema, J. Beyrath, J. Smeitink, B. B. A. de Vries, P.A.C. ’t Hoen, T. Kozicz, N. Nadif Kasri

## Abstract

Mitochondrial encephalomyopathy, lactic acidosis and stroke-like episodes (MELAS) is often caused by an adenine to guanine mutation at m.3243 (m.3243A>G) of the *MT-TL1* gene (tRNA^leu(UUR)^). To understand how this mutation affects the nervous system, we differentiated human induced-pluripotent stem cells (iPSCs) into excitatory neurons with normal (low heteroplasmy) and impaired (high heteroplasmy) mitochondrial function from MELAS patients with the m.3243A>G mutation. We combined micro-electrode array (MEA) measurements with RNA sequencing (MEA-seq) and found that the m.3243A>G mutation affects expression of genes involved in mitochondrial respiration- and presynaptic function, as well as non-cell autonomous processes in co-cultured astrocytes. Finally, we show that the clinical II stage drug sonlicromanol (KH176) improved neuronal network activity in a patient-specific manner when treatment is initiated early in development. This was intricately linked with changes in the neural transcriptome. Overall, MEA-seq is a powerful approach to identify mechanisms underlying the m.3243A>G mutation and to study the effect of pharmacological interventions in iPSC-derived neurons.

**Highlights:**

- High m.3243A>G heteroplasmy leads to lower neuronal network activity and synchronicity
- High heteroplasmy affects expression of genes involved in mitochondrial ATP production and the synaptic function / the presynaptic vesicle cycle
- High neuronal heteroplasmy non cell autonomously affects gene expression in healthy co-cultured astrocytes
- Sonlicromanol partially rescues neuronal network activity and transcriptome changes induced by high heteroplasmy

**eTOC Blurb:** Using human inducible pluripotent stem cell-derived neurons with high levels of m.3243A>G heteroplasmy, Klein Gunnewiek et al. show transcriptome changes underlying the functional neuronal network phenotype, and how sonlicromanol can partially improve both this neuronal network phenotype, and the transcriptome changes, in a patient-specific manner.

## Introduction

Mitochondrial diseases (MD) caused by mutations in the mitochondrial genome (mtDNA), predominantly affects tissues with high energy needs, such as the brain (Andreazza et al., 2018; Gorman et al., 2016; Kim et al., 2019; Pei and Wallace, 2018; Reinhart and Nguyen, 2019; Srivastava et al., 2018; Sullivan et al., 2018; Sylvia et al., 2018). Mitochondrial encephalomyopathy, lactic acidosis and stroke-like episodes (MELAS) is the most common MD. It presents with epilepsy, stroke-like episodes, intellectual and cortical sensory deficits, cognitive decline, muscle weakness, cardiomyopathy, and /or diabetes (El-Hattab et al., 2015). The mtDNA mutation underlying most MELAS cases is an adenine to guanine variant at position m.3243 (m3243A>G) of the *MT-TL1* gene (tRNA^leu(UUR)^), in the mitochondrial genome (mtDNA) (OMIM 590050) (Goto et al., 1990; Hirano and Pavlakis, 1994). Approximately 1:20,000 are clinically affected by this mutation (Chinnery et al., 2000; Hirano and Pavlakis, 1994; Majamaa et al., 1998; Manwaring et al., 2007). Above a threshold mutated mtDNA copies (heteroplasmy percentage) disrupt the normal function of the oxidative phosphorylation system (OXPHOS), with cellular consequences, leading to severe physical clinical signs and symptoms (Ciafaloni et al., 1992; Kobayashi et al., 1990; Schon et al., 2012; Ylikallio and Suomalainen, 2012; Yokota et al., 2015). The m.3243A>G mutation distorts amino acid incorporation during translation of 13 mtDNA proteins, which are part of the OXPHOS subcomplexes I - V. The OXPHOS reduces oxygen to water using electrons from NADH and FADH_2_, producing adenine triphosphate (ATP), and controlling cellular levels of reactive oxygen species (ROS) (Holmström and Finkel, 2014). OXPHOS complex deficiencies lead to a disbalanced cellular redox state, often with increased ROS production (Distelmaier et al., 2009), distorted mitochondrial signaling, and macromolecule damage (lipid peroxidation, etc.) (Daiber, 2010).

Classic mitochondrial disease interventions are focused on the treatment of symptoms (Pfeffer et al., 2012), for instance with antioxidants (coenzyme Q10, riboflavin, or vitamins C and -E (Garrido-Maraver et al., 2012; Glover et al., 2010)), dietary supplements (L-arginine (Parikh et al., 2015)), or increased exercise (Tarnopolsky, 2014). None consistently improve patients’ strength and/or quality of life (Pfeffer et al., 2012). Novel treatments awaiting FDA approval (Lyseng-Williamson, 2016; Rahman and Rahman, 2018) target mitochondrial redox balance, or serve as antioxidants and electron donors (Gorman et al., 2020; Hirano et al., 2018). Sonlicromanol (also known as KH176), full name (S)-6-hydroxy-2,5,7,8-tetramethyl-N-((R)-piperidin-3-yl)chroman-2-carboxamide hydrochloride, is currently in clinical trial stage IIB (Janssen et al., 2019). As a derivative of Trolox, a soluble form of vitamin E, it aims to restore the redox balance, and reduce ROS levels (Beyrath et al., 2018; Koene et al., 2017). This is achieved by modulation of the thioredoxin system / peroxiredoxin enzyme machinery (TrxR-Trx-Prdx system), a three-enzyme chain that reduces H_2_O_2_ into water using electrons from NADPH (Beyrath et al., 2018). Rodent studies detected sonlicromanol in muscle and the brain, in a dose-dependent manner and without marked accumulation, even after frequent daily exposure (Beyrath et al., 2018). Sonlicromanol showed increased life-span and improved performance and gait in Ndufs4^-/-^ mice, with less retinal ganglion cell degeneration (Frambach et al., 2020; Haas et al., 2017). Sonlicromanol’s ROS scavenging abilities, protection from lipid peroxidation, and improved cell viability were shown in complex I deficient human fibroblast lines (Beyrath et al., 2018). Phase I and II clinical trials demonstrated acceptable safety, pharmacokinetic properties (Koene et al., 2017) and tolerance in humans, and improved alertness, as well as reduced depressive symptoms in patients (Janssen et al., 2019).

We have recently established a novel *in vitro* model to study m.3243A>G in human iPSC-derived glutamatergic neurons (iNeurons) (Klein Gunnewiek et al., 2020). iNeurons with high levels of m.3243A>G heteroplasmy (60-65%) had shorter dendrites, less synaptic connections, and reduced and desynchronous network activity. Here, we combined neuronal phenomics (MEA) and transcriptomics (RNA-seq) data to address important research gaps: 1) what mechanisms could be downstream of the impaired neural bioenergetics, and 2) can we reverse neuronal pathology caused by the m.3243A>G variant? Using MEA-seq we confirmed our previous findings (Klein Gunnewiek et al., 2020) on developmental neural network phenotypes, and found reduced expression of genes involved in mitochondrial ATP production and presynaptic function in neurons with high m.3243A>G heteroplasmy. This neuronal phenotype non-cell autonomously induced gene expression changes in healthy co-cultured rat astrocytes. Furthermore, sonlicromanol treatment partially rescued the neuronal network phenotype and gene expression changes in a patient-specific manner.

## Results

### High m.3243A>G heteroplasmy affects neuronal network development

We previously generated iPSC clones, with a wide range of m.3243A>G heteroplasmy levels on an isogenic background, ranging from 0% to 87% (Klein Gunnewiek et al., 2020). We cultured neurons of matching densities (Fig. S1A-C), differentiated from isogenic sets of iPSCs with low levels heteroplasmy (0%; LH1+2) and high levels of heteroplasmy (±60%; HH1+2) as previously described (Klein Gunnewiek et al., 2020). Using micro-electrode array (MEAs) recordings, we observed reduced mean firing rate, increased percentage of random activity, and number of synchronous network bursts, in HH1+2 at 30 days *in vitro* (DIV), consistent with previous findings (Klein Gunnewiek et al., 2020) and reminiscent of less mature neuronal networks (Frega et al., 2017). Other parameters, such as the number of bursts per minute, were not affected (Fig. S1D-I). To exclude the possibility that the neuronal phenotypes observed in the HH lines are due to simple delayed maturation, we recorded network activity at 30, 37, and 43 days in vitro (Fig. 1A). In healthy control networks, the firing- and network burst rates tend to plateau at around DIV28, indicators of a functionally “mature” network (Frega et al., 2017). We found that both, HH1 and HH2 lines, still showed reduced synchronous network bursts up to DIV 43 compared their isogenic LH line (Fig 1B+C). Furthermore, both lines displayed increased percentage of random activity (Fig. 1B+C), and HH1 displayed reduced mean firing rate, at DIV 43.These indicate the HH iNeurons are not simply delayed in development but show a persistent neuronal network phenotype.

**Figure 1.**
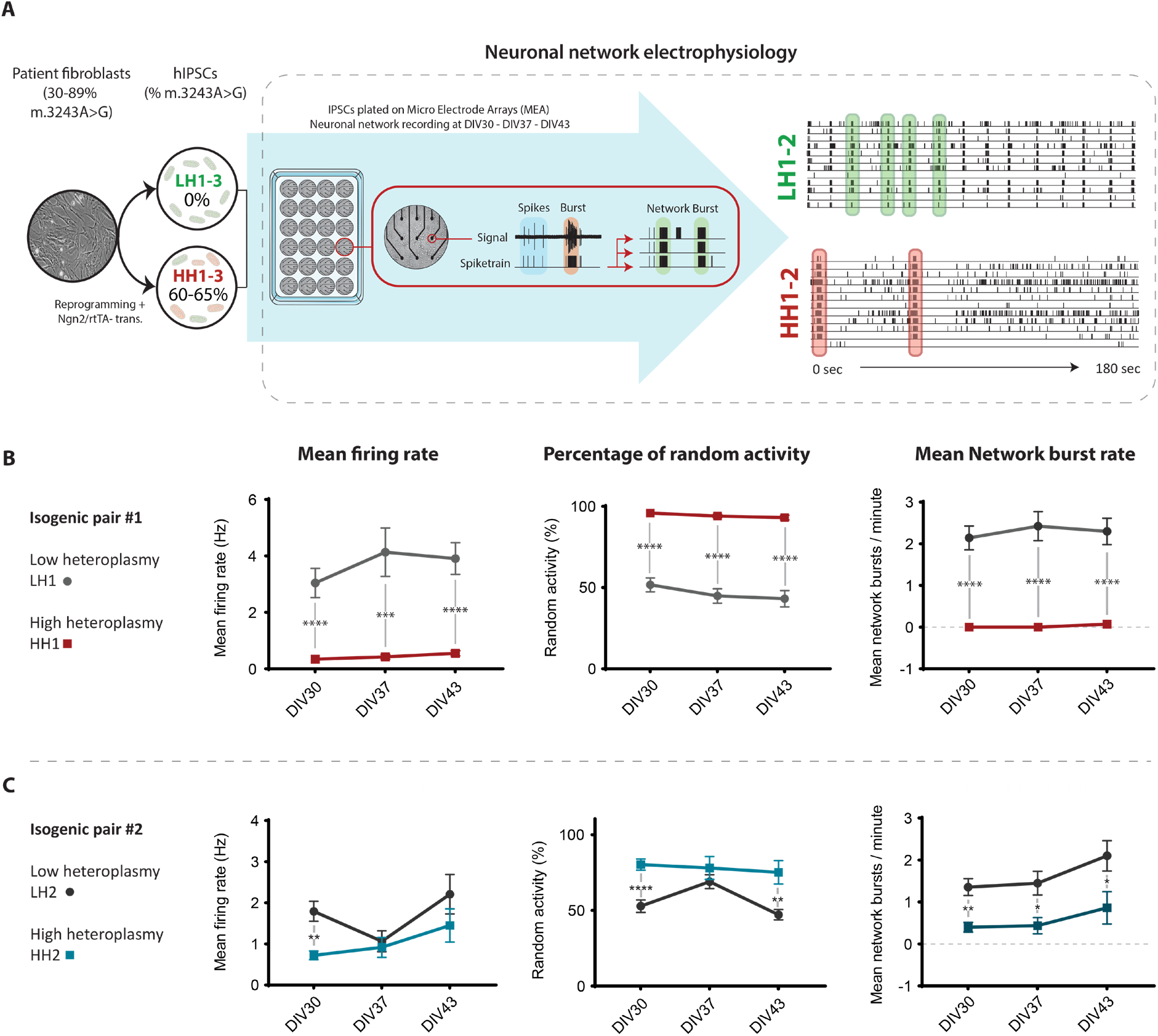
Neuronal network development for LH1+2, and HH1+2 neuronal networks. (A) Patient-derived fibroblasts were reprogrammed to iPSCs, generating low- (0%) and high (60-65%) heteroplasmy clones. These were differentiated into excitatory neurons on micro-electrode arrays (MEAs) recorded for a 10-minute period at 30-, 37-, and 43 DIV. (B-C) Quantification of the mean firing rate, the percentage of random activity, and the network burst rate, for (B) LH1 (n=27) and HH1 (n=27), and (C) LH2 (n=16) and HH2 (n=14). Data represents means ± SEM. *P<0.05, **P<0.01, ***P<0.001, ****P<0.001, using restricted maximum likelihood model, with Holm-Sidak’s correction for multiple comparisons.

### Mitochondrial and synaptic gene expression are affected in m.3243A>G high heteroplasmy iNeurons

To understand the molecular mechanism underlying the neuronal network phenotypes in HH iNeurons we optimized a bulk RNA-seq method which can be used in combination with MEA experiments (MEA-seq). RNA was isolated from iNeurons co-cultured with wild-type (WT) rat astrocytes directly after network activity was measured on DIV 43 (Fig. 2A). After sequencing raw reads were mapped to a combined human- and rat genome to separate reads from iNeurons and rat astrocytes. We confirmed the cell identity of iNeurons and astrocytes, using cell type-specific gene expression (Fig. 2B), i.e. genes known to be highly expressed either in neurons or astrocytes.

**Figure 2.**
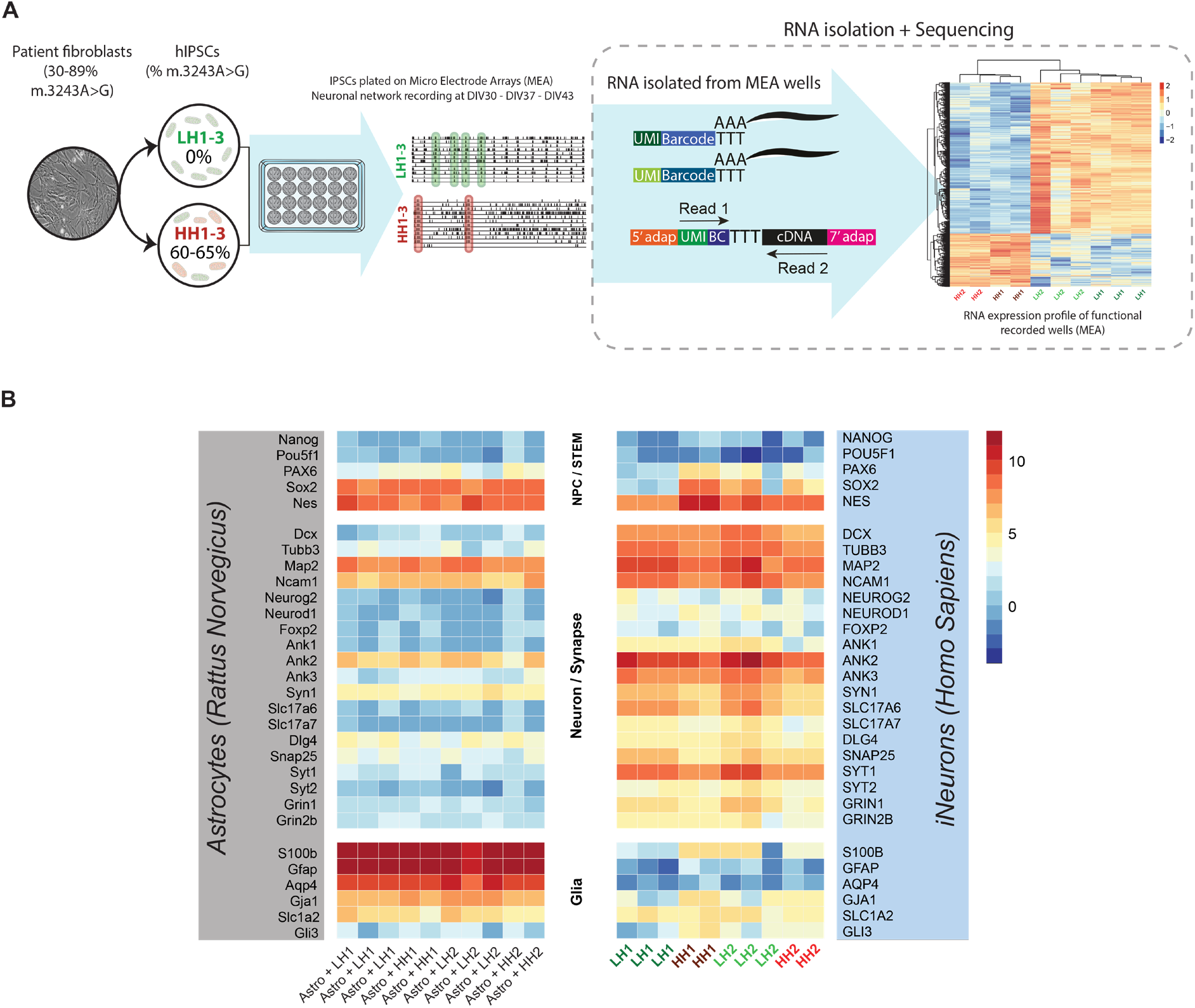
Expression of stem cell, neuronal progenitor cell, neuronal, and glial cell markers. (A) RNA-seq was performed on several representative MEA wells containing iNeurons co-cultured with rat astrocytes. (B) Heatmap showing gene expression levels of stem cell/NPC genes (top section), neuronal- and synaptic genes (middle section), and glial genes (bottom section), including rodent homologs (left section, astrocyte samples), and human homologs (right section, iNeuron samples). Voom-transformed and batch-corrected counts per million (log2 scale) are shown.

Our next aim was to determine how the impaired neuronal bioenergetics caused by high m.3243A>G heteroplasmy levels affect gene expression. Principal component analysis (PCA) showed that HH samples cluster away from LH samples (Fig. S2A). We performed differential expression (DE) analysis followed by gene set enrichment analysis (GSEA) to determine what biological processes were affected in HH iNeurons. Comparing HH samples (HH1 + HH2) with LH samples (isogenic controls, LH1 + LH2), we identified 1580 significantly DE genes (adj. *p* < 0.05); 1169 of which were down-regulated, and 411 of which were up-regulated (Fig. 3A-B, Suppl. table S1). We observed down-regulation of several mtDNA genes (Table S1), probably directly affected by the m.3243A>G mutation as their expression is regulated by MT-TL1. Furthermore, we observed significant down-regulation of nuclear mitochondrial genes, and genes linked to different forms of mitochondrial disease (Thompson et al., 2020) (Table S1). Interestingly, several well-known synaptic genes, as well as several genes linked to epilepsy (Wang et al., 2017), were significantly down-regulated (Table S1), suggesting a potential link with the neuronal network phenotype.

**Figure 3.**
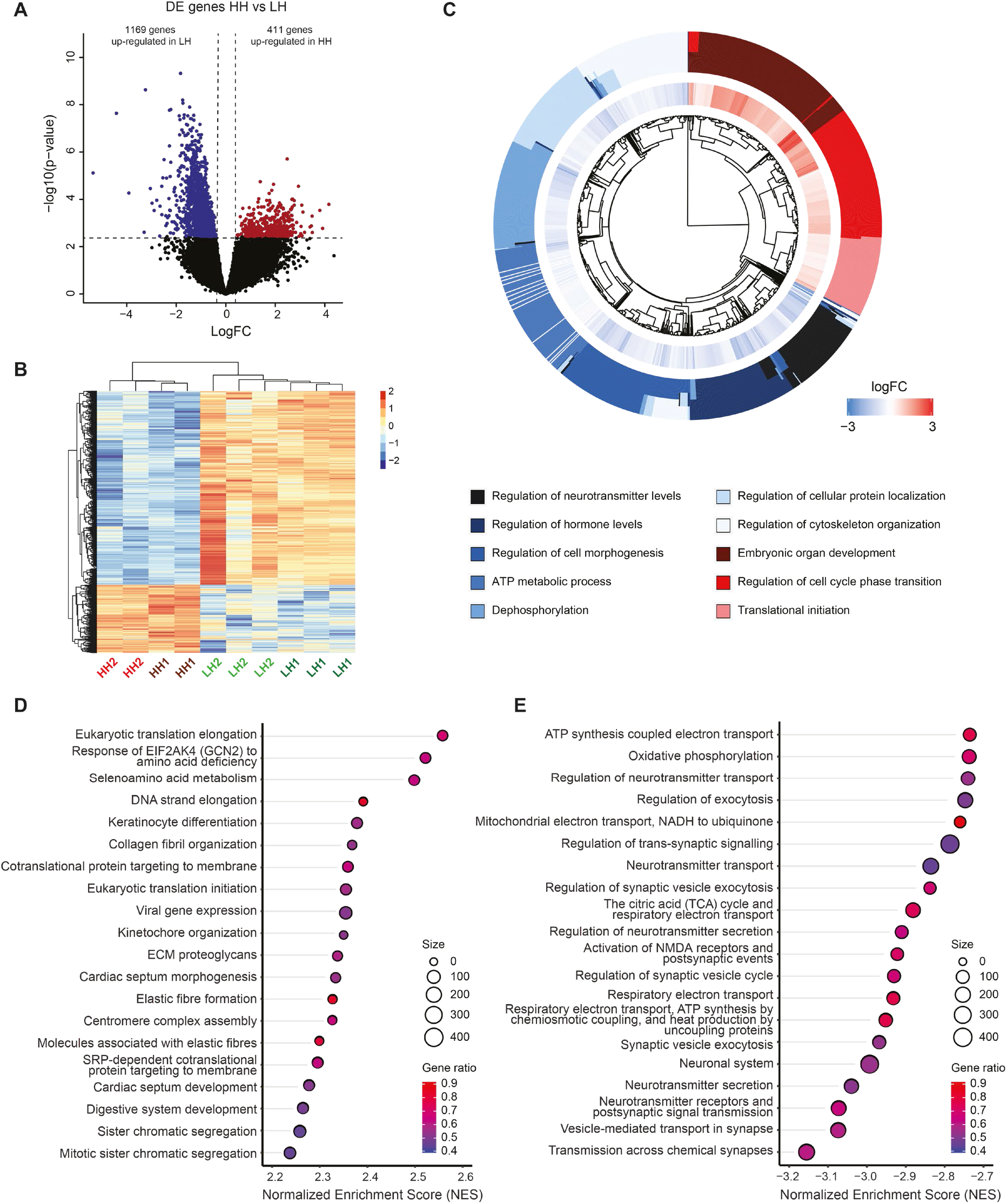
RNA-seq results for HH vs LH iNeurons. (A) Volcano plot showing differentially expressed (DE) genes in HH versus LH iNeurons (adj. *p* < 0.05). (B) Heatmap showing expression of DE genes. Voom-transformed and batch-corrected counts per million (log2 scale) were scaled per gene. (C) Circos plot showing gene sets significantly enriched for up- and down-regulated genes, representing the most important processes affected. LogFC is shown for leading edge genes within each gene set. (D-E) Top 20 gene sets enriched for (D) up-regulated and (E) down-regulated genes. Reactome pathways and GO terms representing biological processes (BP) are shown. Normalized enrichment score (NES) is shown for each gene set, as well as gene set size (circle size) and gene ratio (color gradient).

GSEA revealed enrichment of affected genes in 1717 GO terms and 309 Reactome pathways (adj. *p* < 0.05). Overall, we found changes in key biological processes, such as regulation of neurotransmitter release (i.e. the presynaptic vesicle cycle), cell morphogenesis (i.e. axonal- and neuronal development), and regulation of cell cycle phase transition (Fig. 3C, Suppl. table S1). We observed strongest enrichment of up-regulated genes for processes involved in extracellular matrix structure, DNA strand elongation, centromere complex assembly, and nucleosome DNA binding, amongst others (Fig. 3C), many of which play a role in stem-cell maintenance and cell division. We observed strongest enrichment of down-regulated genes for gene sets and pathways representing (pre)synaptic processes (specifically the synaptic vesicle cycle) and mitochondrial respiration (oxidative phosphorylation, or the use of oxygen in ATP production), as well as neuronal development (Fig. 3D). The RNA-seq data reveal changes in line with the neuronal network phenotype data, showing decreased neuronal activity linked to m.3243A>G heteroplasmy.

### Non-cell autonomous effects of high m.3243A>G heteroplasmy iNeurons on astrocytes

In our model system, neuronal function is critically dependent on co-culturing with rat astrocytes (Frega et al., 2017), as these facilitate synapse formation (Allen and Eroglu, 2017; Allen et al., 2012), synaptic transmission (Witcher et al., 2010), synaptic pruning (Bialas and Stevens, 2013; Chung et al., 2013), and axon formation (Gonçalves et al., 2016). The different origins of cell types (rat versus humans) allowed us to investigate whether neuronal m.3243A>G heteroplasmy level, non-cell autonomously affected gene expression in astrocytes. This is important as astrocytes metabolically support neuronal ATP production by supplying lactate, via the “astrocyte-neuron lactate shuttle” (Araque et al., 1999; Genc et al., 2011; Jimenez-Blasco et al., 2020; Witcher et al., 2010). Gene expression profiles from astrocytes co-cultured with HH iNeurons (Astro+HH1 and Astro+HH2) were compared to astrocytes co-cultured with LH iNeurons (Astro+LH1 and Astro+LH2). Differential expression analysis revealed 79 significant DE genes (adj. *p* < 0.05); 70 up-regulated genes and 9 down-regulated genes (Fig. 4A+B, Fig S2B, Table S1). Interestingly, two down-regulated genes play key roles in mitochondrial ATP production; *Cox4i2* encoding the enzyme that catalyzes the electron transfer from reduced cytochrome c to oxygen; and *Mt-nd1* encoding ETC Complex I subunit (mutations can lead to MELAS (Kirby et al., 2004). GSEA revealed significant enrichment of DE genes in 58 Reactome pathways and 438 GO terms (adj. *p* < 0.05) (Table S1). The top gene sets enriched for up-regulated genes consist of genes involved in extracellular matrix organization, immune response, and cell morphogenesis and differentiation (Fig. 4C). The top gene sets enriched for down-regulated genes contain genes involved in e.g. mitochondrial function, amino acid metabolism, and synaptic function (Fig. 4D). These data show m.3243A>G-induced, non-cell autonomous, changes in neuronal gene expression also reduce astrocytic mitochondrial gene expression, as well as genes involved in synaptic function. How exactly these changes in astrocyte gene expression occur is unclear, but they could result from the neuronal adaptation to their bioenergetic deficit, caused by the high m.3243A>G heteroplasmy.

**Figure 4.**
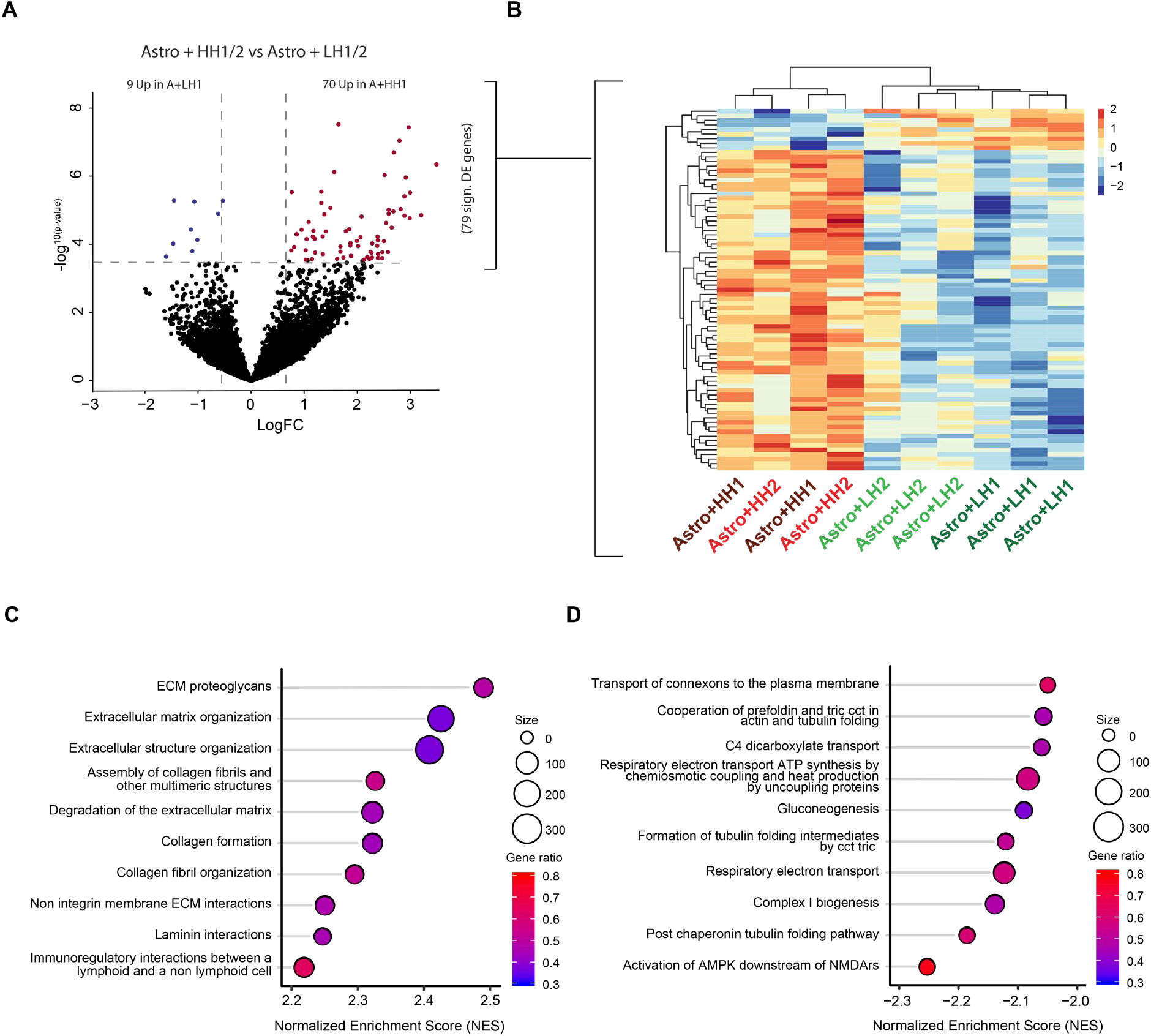
Non-cell autonomous effects of m.3243A>G high heteroplasmy iNeurons on astrocyte gene expression. (A) Volcano plot showing 79 DE genes in astrocytes co-cultured with HH iNeurons compared to astrocytes co-cultured with LH iNeurons. (B) Heatmap showing expression of DE genes. Voom-transformed and batch-corrected counts (log2 scale) were scaled per gene. (C-D) Top 10 gene sets enriched for (C) up-regulated and (D) down-regulated genes are shown. Normalized enrichment score (NES) is shown for each gene set, as well as gene set size (circle size) and gene ratio (color gradient).

### Sonlicromanol improves neuronal network dysfunction of high m.3243A>G heteroplasmy iNeurons in a patient-specific manner

We applied the ROS scavenging- and redox state modulator sonlicromanol (KH176) (Beyrath et al., 2018; Haas et al., 2017; Koene et al., 2017) to young (DIV 3)- and mature (DIV 29) HH neuronal networks, to test its ability to rescue the HH neuronal network phenotypes. We treated MEA-grown neuronal networks short-term (two weeks; DIV 29-DIV 43) and long-term (six weeks; DIV 3-DIV 43), with different concentrations of sonlicromanol (0.5 μM, 1 μM, 3 μM and 5 μM + DMSO vehicle condition), and recorded network activity at 30, 37, and 43 DIV (Fig. 5A-G). We chose these concentrations based on work by Beyrath et al., (2018), who observed improved cell viability at 0.1 – 1 μM sonlicromanol, with an EC50 = 0.27 μM, in fibroblasts with mutations in different nuclear encoded Complex I subunits.

**Figure 5.**
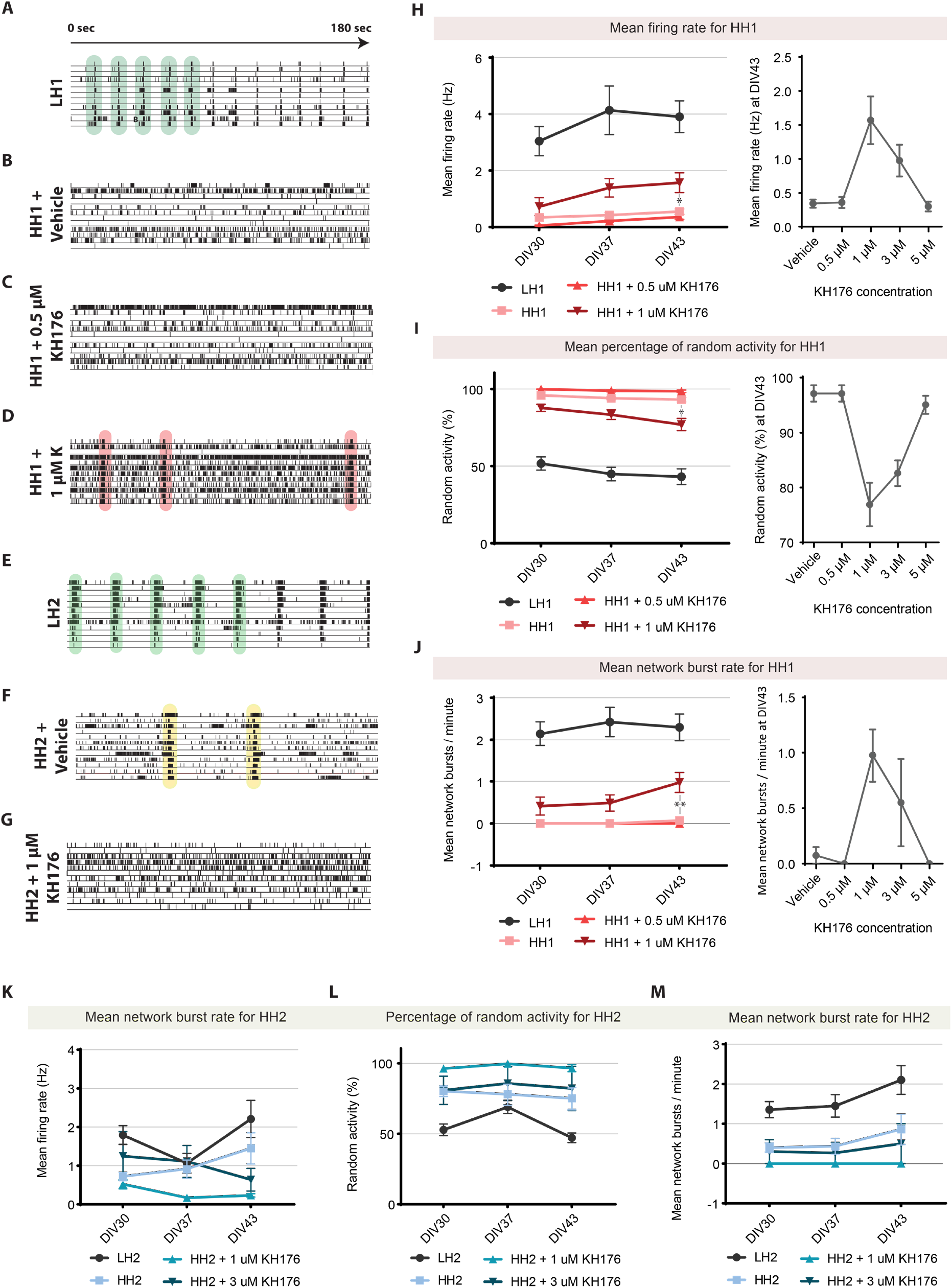
Neuronal network activity after sonlicromanol (KH176) treatment starting from 3 days in vitro, in HH1 and HH2 iNeuronal networks. (A-G) Representative raster plots of 3 minute neuronal network activity recordings at DIV 43 of indicated conditions (A) LH1 (n = 17), (B) HH1 (n = 12), (C) HH1 + 0.5 μM sonlicromanol (n = 8), (D) HH1 + 1 μM sonlicromanol (n = 8), (E) LH2 (n = 6), (F) HH2 (n = 16), and (G) HH2 + 1 μM sonlicromanol (n = 4). (H) Quantification of the overall mean firing rate, (I) Quantification of the percentage of random activity, (J) Quantification of the network bursts rate. Data represents means ± SEM. *P<0.05, **P<0.01, ***P<0.001, ****P<0.001, using restricted maximum likelihood model, with Holm-Sidak’s correction for multiple comparisons between treated and untreated samples.

Short-term treatment of a mature network from 29 DIV onwards, had no positive effect on any of the key MEA parameters, in either patient line, at either concentration (Fig. S3). When we treated the neuronal networks (HH1 + HH2) from an early point in neuronal development (from 3 DIV onwards), we observed patient-specific improvements in MEA response. Specifically, 1 μM sonlicromanol most significantly improved MEAs parameters in HH1. We found a significant increase in mean firing rate (*p*<0.01) and network burst rate (*p*<0.001), and decreased percentage of random activity (*p*<0.005), compared to the untreated HH1 or HH1-vehicle conditions (Fig. 5H-J). We observed similar, albeit less pronounced improvements at a concentration of 3 μM (Fig. 5H-J). No reversal of pathology was observed using 5 μM sonlicromanol, suggesting that the effects of sonlicromanol is dose dependent. Of note, the improvements seen in HH1 with sonlicromanol treatment at 1 μM and 3 μM did not generalize to line HH2 (Fig. 5K-M).

Together, this shows that for patient HH1, sonlicromanol could be beneficial in restoring some of its neuronal network activity and synchronicity, when treatment is initiated early in neuronal network development, as opposed to treatment at a later stage. However, these improvements are mild, dose-dependent, and did not translate to a second patient line.

### Sonlicromanol reverses changes in gene expression in high m.3243A>G heteroplasmy iNeurons in a patient-specific manner

To determine the effect of the sonlicromanol treatment on gene expression level, we isolated RNA directly from MEA wells with a representative neuronal phenotype, and performed DE analysis comparing sonlicromanol treated versus untreated samples (Fig. S4A; Table S2). As only HH1 showed functional improvement on MEA following sonlicromanol treatment, we expected patient-specific drug responses on gene expression level as well, therefore comparisons were made for each isogenic set separately. PCA showed HH1+KH176 (sonlicromanol) samples clustered away from untreated HH1 samples, towards the direction of LH1 samples, indicating sonlicromanol had positive effects on gene expression (Fig. S2C). Differential expression analysis for the first isogenic set (HH1 vs LH1) revealed 1715 significantly DE genes (adj. *p* < 0.05); 846 down-regulated and 869 up-regulated, in HH1 samples (Fig. 6A, Table S2). Sonlicromanol treatment of the HH1 line (HH1+KH176) compared to untreated HH1 revealed 116 DE genes (adj. *p* < 0.05); 112 down-regulated and 4 up-regulated genes (Fig. 6B, Table S2). Interestingly, out of these 116 DE genes, 113 genes overlapped with genes differentially expressed in HH1 compared to LH1 (Fig. 6D). Hierarchical clustering of samples on HH1 vs LH1 DE genes reveals sonlicromanol treated HH1 samples clearly cluster together with LH1 (Fig. 6C), showing the sonlicromanol treatment directed the expression of genes in HH1 towards control level. GSEA revealed enrichment of affected genes in 1869 GO terms and 310 Reactome pathways (adj. *p* < 0.05) for the HH1 vs LH1 comparison (Fig. 6E, Table S2), and identified the same biological processes as were affected in both HH lines (Fig. 3C-E). When we performed GSEA for the sonlicromanol treated versus untreated HH1 samples, we found enrichment of affected genes in 1439 Go terms and 243 Reactome pathways (Table S2). We observed strong enrichment for down-regulated genes in gene sets involved in different aspects of cell division, from nuclear division, to pattern specification, and chromatid segregation, which matched the processes that were up-regulated in HH1 compared to LH1. Also, strongest enrichment of up-regulated genes was observed for gene sets that were shown to be most strongly enriched for down-regulated genes linked to m.3243A>G heteroplasmy; which included gene sets involved in mitochondrial respiration and (pre)synaptic function (Fig. 6F+G+H, Fig. S4B-D). The increased expression of synaptic genes due to sonlicromanol treatment supported the findings from neuronal network activity measurements showing functional improvements in sonlicromanol treated HH1 iNeurons.

**Figure 6.**
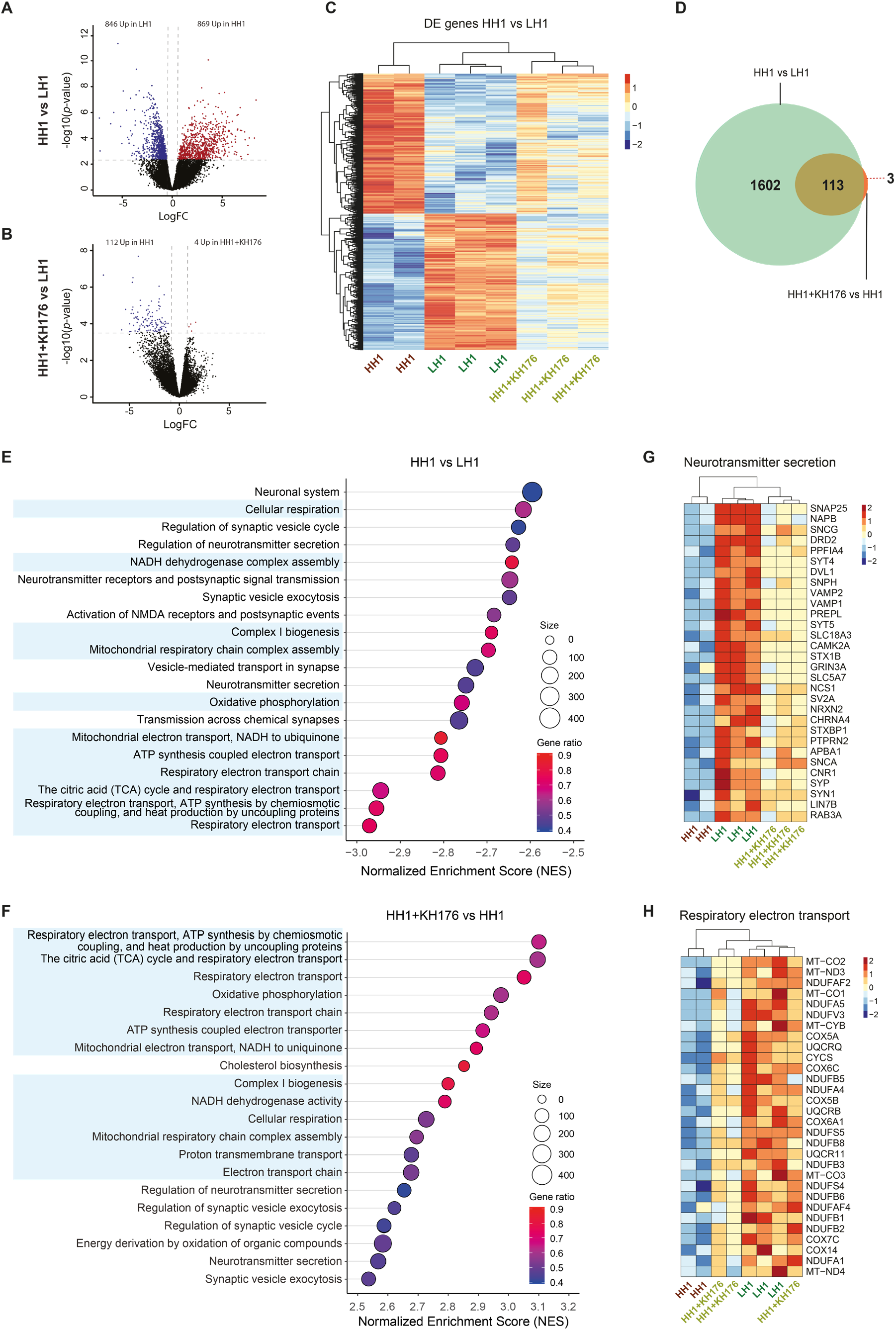
Effects of sonlicromanol (KH176) treatment on gene expression of m.3243A>G high heteroplasmy iNeurons of isogenic set 1 (HH1). (A) Volcano plot showing 1715 DE genes (adj. *p* < 0.05) in HH1 compared to LH1 samples. (B) Volcano plot showing 116 DE genes (adj. *p* < 0.05) in HH1+KH176 compared to HH1 untreated. (C) Heatmap showing expression of DE genes in HH1 vs LH1, for LH1, HH1 and HH1+KH176 samples. Voom-transformed and batch-corrected counts per million (log2 scale) were scaled per gene. (D) Venn diagram showing overlap between DE genes in HH1 vs LH1, and DE genes in HH1+KH176 vs HH1. (E-F) Top 20 gene sets enriched for down-regulated genes in (E) HH1 vs LH1 and (F) HH1+KH176 vs HH1. Reactome pathways and GO terms representing biological processes (BP) are shown. Normalized enrichment score (NES) is shown for each gene set, as well as gene set size (circle size) and gene ratio (color gradient). Gene sets that represent processes involved in mitochondrial ATP production are marked blue. (G-H) Heatmap showing top 30 overlapping leading edge genes (both for HH1 vs LH1, and for HH1+KH176 vs HH1) for (G) the neurotransmitter secretion gene set and (H) the respiratory electron transport gene set. Voom-transformed and batch-corrected counts were scaled per gene.

Next, we analyzed the RNA-seq data for the second isogenic set separately. We first compared gene expression profiles from HH2 to LH2. Differential expression analysis revealed 857 DE genes (adj. *p* < 0.05), of which 197 are up-regulated and 660 are down-regulated (Fig. S5A+B, Table S2). Interestingly, no significant DE genes were identified when comparing sonlicromanol treated to untreated HH2 samples. A heatmap of HH2 vs LH2 DE genes show HH2 samples treated with sonlicromanol cluster together with HH2, whereas LH2 samples cluster away, confirming the compound has minor to no effects on expression of genes affected in HH2 (Fig. S5B, Fig. S2D), in line with our results from neuronal network activity measurements.

## Discussion

In this study, we integrated neuronal phenomics with transcriptomics data to provide novel insight into potential mechanisms underlying changes in neuronal network activity linked to high levels (60%) of m.3243A>G heteroplasmy. Specifically, we observed down-regulation of both mitochondrial- and nuclear genes involved in mitochondrial ATP production, matching the reduced mitochondrial function previously observed in human neurons with m.3243A>G heteroplasmy (Klein Gunnewiek et al., 2020). Furthermore, genes involved in (pre)synaptic processes were down-regulated, in line with decreased neuronal network activity, and previously observed single cell presynaptic deficits (Klein Gunnewiek et al., 2020). Finally, we provide encouraging data for sonlicromanol as a potential novel treatment for MELAS. Specifically, sonlicromanol treatment starting from an early point in neuronal network development resulted in the up-regulation of genes involved in mitochondrial respiration and (pre)synaptic function in HH1 samples, as well as improved neuronal network activity, due to sonlicromanol treatment starting from an early point in development.

Consistent synaptic function depends in part on mitochondrial ATP production (Rangaraju et al., 2014). The presynaptic vesicle cycle requires most presynaptic ATP, and is negatively affected by dysfunctional ATP synthesis (Rangaraju et al., 2014). We previously found less mitochondrial oxygen consumption and increased glycolysis in HH iNeurons, with less Synapsin 1/2 puncta and less mitochondria at presynaptic sites (Klein Gunnewiek et al., 2020). Here, we found down-regulation of mtDNA genes encoding for subunits of OXPHOS complexes I, III, IV, and V, and several nuclear genes coding for Complex I (Table S1). Expression of genes encoding for Complex II subunits were unaffected, as were most nuclear genes encoding for Complex III, IV, and V. We did observe down regulation of other genes linked to mitochondrial disease (Rahman and Rahman, 2018; Thompson et al., 2020), involved in OXPHOS, pyruvate transport, ketone body catabolism, mitochondrial trafficking, cleavage of membrane phospholipids, and other functions (Table S1). Furthermore, we also observed reduced expression of *SYN1* and *SYN2* (Synapsin 1/2), as well as other synaptic genes such as GRIA4, GRIN1/3A, or VAMP1/2 (Table S1), in which mutations have been linked to for instance Alzheimer’s disease (Berchtold et al., 2013). Down regulation of *SYT1/4/5/6/7*, which encode for the presynaptic Ca^2+^ sensor Synaptotagmins, point to distorted presynaptic Ca^2+^ homeostasis, which affects synaptic release probabilities (Kwon et al., 2016), and could partially explain synaptic deficits we observed here, and previously (Klein Gunnewiek et al., 2020). Reduced *PINK1* expression also stands out, as *PINK1* mutations are linked to reduced dendritic growth (Dagda et al., 2014), reduced Complex I- and synaptic function (Morais et al., 2009), irregular Parkin-mediated mitophagy (Dagda et al., 2009), and Parkinson’s disease (Agnihotri et al., 2017; Lee et al., 2017; Verma et al., 2018). Epilepsy is a frequent presentation of MELAS. Although epilepsy is more often linked to inhibitory neuron dysfunction (Kann, 2016; Magloire et al., 2019; Patel et al., 2019), here we did observe down-regulation of genes directly linked to epilepsy, in excitatory neurons (Wang et al., 2017), such as *STXBP1* (Table S1). Interestingly, loss of function mutations in *STXBP1* are linked to infantile epileptic encephalopathy and non-syndromic epilepsy (Pavone et al., 2012). *STXBP1* knock-down also reduces sEPSC frequency (not amplitude) in human iNeurons (Zhang et al., 2013), similar to what we observed previously in HH iNeurons (Klein Gunnewiek et al., 2020).

We report that sonlicromanol, a compound targeting mitochondrial function, can improve both neuronal network function as well as changes in gene expression, representing mitochondrial respiration and synaptic function. Sonlicromanol has not yet been tested in a large-scale clinical trial, but the phase II clinical trial has already shown improvements in mood and alertness, often found in patients suffering from m.3243A>G related- and other mitochondrial diseases (Janssen et al., 2019). Our findings provide molecular and functional evidence for potential positive effects of sonlicromanol on these neuronal specific deficits. Specifically, genes that were affected by m.3243A>G heteroplasmy, were reversely affected by sonlicromanol treatment in patient HH1.The improved neuronal function seen on MEA was reflected by up-regulation of genes involved in presynaptic function. We noted no improvements in HH2 following sonlicromanol treatment. The patient-specific improvements we observe, both on a functional level (MEA) as well as on a transcriptome level, suggest that personalized testing like “N=1” clinical trials (Schork, 2015), might be more suited for MELAS and mitochondrial disease, where average performance scores could blur promising findings. Additionally, whereas both have similar heteroplasmy levels, and the same processes affected in HH1 are affected in HH2. It is unclear why only one HH line responded to the sonlicromanol treatment, however, it could represent the large phenotypic variability in patients with m.3243A>G related mitochondrial disease. Future work using MELAS iPSC lines generated from responders and nonresponders, could predict treatment response.

As a final note, we found astrocyte gene expression was affected by the presence of iNeurons with high m.3243A>G heteroplasmy levels. Genes involved in metabolism and tubulin formation were downregulated in astrocytes co-cultured with HH iNeurons. Neuronal activity facilitates astrocyte maturation and expression of astrocyte genes with a role for Notch signaling (Hasel et al.; 2017). Both neuronal Notch ligands and astrocyte Notch-1/2 receptors are needed for astrocyte glutamate uptake (Hasel et al., 2017), and whilst *Notch1/2* expression is unaffected in astrocytes co-cultured with HH iNeurons, the HH iNeurons do show reduced expression of Notch ligands *JAG2* and *DLK2*. This impaired balance could further add to the observed synaptic phenotype and could represent a novel treatment target.

The integration of phenomics and transcriptomics data have disentangled several neuronal features of MELAS, advancing the understanding of the impact of m.3243A>G heteroplasmy on the human nervous system. Our findings on neuronal responses to sonlicromanol treatment confirm the heterogeneity and individualized nature of patient response to treatment, as well as the need for treatments tailored to the needs of the individual mitochondrial disease.

## Experimental Procedures

### iPSC generation and culture

These methods have been described in detail previously (Klein Gunnewiek et al., 2020). In short, fibroblasts of a MELAS subject with the pathogenic variant m.3243A>G in *MT-TL1* (tRNA^Leu(UUR)^) were reprogrammed through retroviral transduction of the Yamanaka transcription factors Oct4, c-Myc, Sox2 and Klf4 (Takahashi and Yamanaka, 2006). Normal karyotypes were confirmed for one iPS clone with 0% m.3243A>G heteroplasmy (“Low heteroplasmy 1”; LH1), and one iPS clone with 65% m.3243A>G heteroplasmy (“High heteroplasmy 1”; HH1). Lines LH2 and HH2 were generous gifts from Ester Perales-Clemente and Timothy Nelson, previously characterized and tested for heteroplasmy levels (Table S1), and with normal karyotypes (Perales-Clemente et al., 2016). iPSCs for subject #2 (LH2 and HH2) were originally reprogrammed from fibroblasts using CytoTune-iPS 2.0 Sendai Reprogramming (Invitrogen A16517). All iPSCs were kept under 15 passages after initial heteroplasmy measurement, to ensure heteroplasmy levels are as previously measured, with medium changes every 2-3 days and passaging 1-2 times per week.

### Neuronal differentiation

iPSCs were derived into upper-layer, excitatory cortical neurons by overexpressing Neurogenin 2 (Ngn2), as described previously. rtTA/Ngn2-positive iPSCs were plated as single cells at DIV0 onto 24-well multi-electrode arrays (Multichannel Systems, MCS GmbH, Reutlingen, Germany), coated with 50μg/mL poly-L-ornithine hydrobromide (PLO; Sigma-Aldrich #P3655-10MG) and 5 μg/mL human recombinant laminin 521 (BioLamina #LN521-02) in E8 basal medium (Gibco #A1517001) supplemented with 1% Penicillin/Streptomycin (Pen/Strep; Sigma-Aldrich P4333), 1% RevitaCell (Thermo-Fisher #A2644501), and 4 μg/mL doxycycline (Sigma-Aldrich #D9891-5G) to drive Ngn2 expression, at 20,000 cells per well for LH1+2, and 30,000 cells per well for HH1+2, to ensure similar mature neuron cell density. At DIV1, medium was changed to DMEM/F12 (Gibco #11320-074) supplemented with 1% Pen/Strep, 4 μg/mL doxycycline, 1% N-2 supplement (Gibco #17502-048), 1% MEM non-essential amino acid solution (NEAA; Sigma-Aldrich #M7145), 10 ng/mL human recombinant BDNF (Promokine #C66212), 10 ng/mL human recombinant NT-3 (Promokine #C66425). To support the neuronal culture, freshly prepared primary cortical rat astrocytes (isolated as previously described by Frega et al., 2017) were added on DIV2, in a 1:1 ratio. At DIV3, the medium was changed to Neurobasal (Gibco #21103-049), supplemented with 20 μg/mL B-27 (Gibco #0080085SA), 1% GlutaMAX (Gibco), 1% Pen/Strep, 4 μg/mL doxycycline, 10 ng/mL human recombinant NT3, 10 ng/mL human recombinant BDNF. At DIV3 only, 2 μM Cytosine β-D-arabinofuranoside (Ara-C; Sigma-Aldrich C1768-100MG) was added to remove any proliferating cells. From DIV5-DIV43 50% of the medium was refreshed every 2 days, supplemented with 2.5% Fetal Bovine Serum (FBS; Sigma-Aldrich #F2442-500ML) from DIV9 onwards.

### Micro-electrode array recordings

Recordings of the spontaneous activity of neuronal networks derived from four iPSC lines (LH1+2, HH1+2) were performed at DIV 30, 37, and 43. All recordings were performed using the 24-well MEA system (Multichannel Systems, MCS GmbH, Reutlingen, Germany). After a 10-minute acclimatization period (37°C; 5% CO2), spontaneous neuronal network activity was recorded for a subsequent 10 minutes. The signal was sampled at 10 KHz, filtered with a high-pass filter (i.e. Butterworth, 100 Hz cut-off frequency) and the noise threshold was set at ± 4.5 standard deviations. Data analysis was performed off-line by using Multiwell Analyzer (i.e. software from the 24-well MEA system that allows the extraction of the spike trains) and a custom software package named SPYCODE developed in MATLAB (The Mathworks, Natick, MA, USA) that allows the extraction of parameters describing the network activity (Bologna et al., 2010), as previously described (Klein Gunnewiek et al., 2020). Thresholds to determine the mean firing rate (MFR), the mean burst rate (MBR), and the mean network burst rate were set as previously described (Klein Gunnewiek et al., 2020).

Sonlicromanol (KH176), full name (S)-6-hydroxy-2,5,7,8-tetramethyl-N-((R)-piperidin-3-yl)chroman-2-carboxamide hydrochloride, was provided by Khondrion as a powder for reconstitution in dimethyl sulfoxide (DMSO) (Beyrath et al., 2018; Koene et al., 2017). DMSO was used as vehicle. Sonlicromanol was added at 100 nM, 300 nM, 500 nM, 1μM, 3 μM, or 5 μM, from either DIV3 (early in development), or from DIV29 (mature neuronal network), up to DIV43. To maintain drug levels in the culture medium, it was added during every medium change.

### RNA sequencing

RNA-seq was performed on hiPSC-derived neurons from subject #1 and #2, including low heteroplasmy (LH) samples, high heteroplasmy (HH) samples, and HH samples treated with sonlicromanol. RNA was isolated after measuring network activity of the neurons on MEAs at DIV43, from 2-3 biological replicates per condition. LH and HH samples were distributed across two MEA batches. HH samples treated with sonlicromanol were included only in the second batch. RNA was isolated with the Quick-RNA Microprep kit (Zymo Research, R1051) according to manufacturer’s instructions. RNA quality was checked using Agilent’s Tapestation system (RNA High Sensitivity ScreenTape and Reagents, 5067-5579/80). RIN values ranged between 6.4 – 9.1. Library preparation was performed using a published single-cell RNA-seq protocol (Cao et al., 2017), adapted for bulk RNA-seq experiments.

For each sample 25 ng total RNA was used as input for RNA-seq library preparation. In short, an anchored oligo-dT primer was used for reverse transcription, followed by second strand synthesis and subsequent removal of excess primers using Exonuclease I (NEB, M0293). cDNA samples were pooled per sets of 8, randomized across three pools, and a 1.2x Ampure XP Beads clean-up was performed (Beckman Coulter, A63881). Next, tagmentation was performed using TDE1 Enzyme (Illumina, 15027865), followed by a 2.0x beads clean-up. PCR amplification was performed for 15 cycli using the NEBNext High-Fidelity 2X PCR Master Mix (NEB, M0541), followed by a 0.8x beads clean-up. Gel extraction was performed to select for products between 200-1000 bp. cDNA concentrations of the final libraries were measured by Qubit dsDNA HS Assay kit (Invitrogen, Q32854). Product size distributions were visualized using Agilent’s Tapestation system (D5000 ScreenTape and Reagents, 5067-5588/9). Libraries were sequenced on the NextSeq 500 platform (Illumina) using a V2 75 cycle kit (Read 1: 18 cycles, Read 2: 52 cycles, Index 1: 10 cycles). A full description of the RNA-seq library preparation can be found in the Supplemental Material.

### Pre-processing of RNA-seq data

Base calls were converted to fastq format and demultiplexed using Illumina’s bcl2fastq conversion software (v2.16.0.10) tolerating one mismatch per library barcode. Reads were filtered for valid unique molecular identifier (UMI) and sample barcode, tolerating one mismatch per barcode. Trimming of adapter sequences and over-represented sequences was performed using Trimmomatic (version 0.33) (Bolger et al., 2014). Trimmed reads were mapped to a combined human (GRCh38.p12) and rat (Rnor_6.0) reference genome to separate reads belonging to the human iNeurons from reads originating from the rat astrocytes. Mapping was performed using STAR (Dobin et al., 2013) (version 2.5.1b) with default settings (--runThreadN 1, --outReadsUnmapped None, --outFilterType Normal, --outFilterScoreMin 0, --outFilterMultimapNmax 10, --outFilterMismatchNmax 10, --alignIntronMin 21, --alignIntronMax 0, --alignMatesGapMax 0, --alignSJoverhangMin 5, --alignSJDBoverhangMin 3, --sjdbOverhang 100). Uniquely mapped reads (mapping quality of 255) were extracted and read duplicates were removed using the UMI-tools software package (Smith et al., 2017). Raw reads from BAM files were further processed to generate count matrices with HTSeq (Anders et al., 2015) (version 0.9.1) using a combined Gencode GRCh38.p12 (release 29, Ensembl 94) and rat (Rnor_6.0) reference transcriptome to count reads mapping to the human and rat genome, belonging to human iNeurons and rat astrocytes, respectively.

### RNA-seq data analysis

Raw counts from count tables were transformed to counts per million (cpm) using edgeR version 3.26.8 (R package) (Robinson et al., 2009). Transcripts with a cpm > 2 in at least two samples were included. Counts were voom-transformed (log2-transformation on cpm values), and corrected for MEA batch effect and genetic background for differential expression (DE) analysis using limma version 3.40.6 (R package) (Ritchie et al., 2015). A linear regression model was fit, in which the voom-transformed expression values were modelled as a function of the condition (LH untreated, HH untreated, HH treated), the isogenic set (set#1, set#2), and MEA batch (plate 1, plate 2). For DE analysis between HH and LH lines, a contrast was defined comparing HH untreated with LH untreated samples using the following design formula: model.matrix(^~^0+condition+set+MEA). DE analysis on RNA-seq data from astrocytes co-cultured with HH and LH iNeurons was performed using the same approach. For DE analysis between HH treated and HH untreated samples, pairwise comparisons were made within each isogenic set, using the following design formula: model.matrix(^~^0+condition+MEA). Genes with a Benjamini-Hochberg (BH)-corrected *p*-value < 0.05 were considered to be significantly differentially expressed between two conditions.

Gene set enrichment analysis (GSEA) was performed using fgsea version 1.10.1 (R package) (Korotkevich and Sukhov, 2019). Genes were ranked based on the *t*-statistic from the DE analysis. Enrichment of genes was tested in Gene Ontology (GO) terms (C5 collection), Reactome pathways (C2 canonical pathways sub-collection) and in NRF2 transcription factor target (TFT) gene sets (C3 TFT sub-collection) from the Molecular Signatures Database (MSigDB) using msigdbr version 7.1.1 (R package) (Igor Dolgalev, 2019). Gene sets were loaded for the correct species (Homo Sapiens or Rattus Norvegicus) per enrichment analysis (iNeuron or rat astrocyte samples, respectively). Gene symbols corresponding to transcripts not part of the final gene list were removed from the selected gene sets. Subsequently, gene sets with remaining gene set size > 5 and < 500 were used for GSEA. Gene symbols (Ensembl version 94) from the final gene list were converted to gene symbols from Ensembl version 97, corresponding to the version of gene symbols used in MSigDB. Gene sets with a Benjamini-Hochberg (BH)-corrected *p*-value < 0.05 were considered to be significantly enriched for up- or down-regulated genes.

### RNA-seq data visualization

Principal component analysis (PCA) was performed on the voom-transformed and batch corrected counts using the prcomp function from stats version 3.6.1 (R package). Volcano plots were generated by plotting the logFC against the −log10 *p*-values from DE analysis results. Heatmaps were generated using voom-transformed and batch corrected counts scaled per gene, with hierarchical clustering performed on rows and columns. Results from GSEA were visualized by plotting the normalized enrichment scores (NES), gene set size and gene ratio for the top 10-20 gene sets based on highest NES, for either up- or down-regulated genes. The gene ratio is calculated by dividing the number of leading edge genes by gene set size. Also, a Circos plot was generated using the GOCluster function from GOplot version 1.0.2 (R package), which shows the logFC for the leading edge genes per gene set. Enrichment plots were generated using the plot Enrichment function using Rcpp version 1.0.2 (R package) (Eddelbuettel and Balamuta, 2018).

### Statistical analysis

Analysis was done using unpaired t-tests, one-way analysis of variance with Bonferroni post hoc correction, or one-way repeated measures ANOVA with sequential post hoc Bonferroni corrections, or Kolmogorov-Smirnov test, where appropriate, using GraphPad Prism 6 (GraphPad Software). *p*-values of *p*<0.05 and smaller, were deemed significant. Sample sizes were based on our previous experiences in the calculation of experimental variability. The output of all analyses is grouped per figure and combined in supplementary table S1.

## Acknowledgements

The raw data have been deposited with the Gene Expression Omnibus (www.ncbi.nlm.nih.gov/geo) under accession code GSE154825. We are grateful to Eva Morava and David Cassiman of the University Hospital Leuven, and Ester Perales-Clemente and Timothy Nelson (Mayo Clinic, RC, MN) for their generous donation of MELAS iPSCs. We thank the department of Molecular Developmental Biology at the Radboud Institute of Molecular Life Sciences for RNA library sequencing. We thank Khondrion for permission to use the sonlicromanol. This work was made possible by the generosity of the Marriott family (to T.K.) and supported by the Tjalling Roorda Foundation (to T.M.K.G.), Stichting Stofwisselingskracht (project number 2017-20 to T.K. and N.N.K.), Netherlands Organization for Health Research and Development ZonMw grant 91217055 (to N.N.K.), ERA-NET NEURON DECODE! grant (NWO) 013.18.001 (to N.N.K.), and Epilepsiefonds WAR 18-02 (to N.N.K.).

## Author contributions

T.M.K.G. and A.H.A.V. performed the experiments and analyzed the data. T.M.K.G., A.H.A.V., T.K., and N.N.K. conceived the hypothesis and designed the experiments. M.H., and C.S. assisted in experiments and technical optimization. H.R., J.B., and J.S. provided sonlicromanol and assistance. T.M.K.G. and A.H.A.V. drafted the manuscript. All authors edited the draft manuscript.

## Supplementary data

### Supplementary methods

**Table S1. DE + GSEA analysis of iNeurons HH1+2 vs LH1+2; Table of various genes of interest; DE + GSEA analysis of Astrocyte genes HH1+2 vs LH1+2 (Excel file)**

**Table S2. DE + GSEA analysis of iNeurons HH1 vs LH1, including KH176 treatment; DE + GSEA analysis of iNeurons HH2 vs LH2, including KH176 treatment (Excel file).**

## Supplementary methods

### RNA-seq library preparation

For each sample, 25 ng total RNA (in 0.65 μL) was mixed with 0.1 μL dNTP mix (10 mM each) (Invitrogen, 10297018), 0.3 μL nuclease-free water (NF H2O) and 0.4 μL anchored oligo-dT (2.5 μM) primer (5’-ACGACGCTCTTCCGATCTNNNNNNNN[10bp index]TTTTTTTTTTTTTTTTTTTTTTTTTTTTTTVN-3’, where “N” is any base and “V” is either “A”, “C” or “G”; IDT) in a tube containing 7 μL Vapor-Lock (Qiagen, 981611). Each sample was incubated for 5 min at 65°C and directly placed on ice. First strand reaction mix was added, consisting of 0.4 μL Maxima RT buffer (5X) (Thermo Scientific, EP0751), 0.05 μL RNasin Plus (Promega, N2611) and 0.1 μL Maxima H Minus Reverse Transcriptase (Thermo Scientific, EP0751). Reverse transcription was performed by incubation at 50°C for 30 min and terminated by heating at 85°C for 5 min. 2 μL RT product was mixed with 7.7 μL NF H2O, 2.5 μL Second Strand Buffer (Invitrogen, 10812014), 0.25 μL dNTP mix (10 mM each), 0.35 μL DNA polymerase I (E. coli) (NEB, M0209), 0.09 μL DNA ligase (E. coli) (NEB, M0205) and 0.09 μL Ambion RNase H (E. coli) (Invitrogen, AM2293). Second strand synthesis was performed by incubation at 16°C for 150 min, followed by 75°C for 20 min. 0.5 μL Exonuclease I (NEB, M0293) was added per sample and incubated at 37°C for 60 min. cDNA samples were pooled per sets of 8, randomized across three pools. Vapor-Lock was removed, and samples were added up with NF H2O to a total volume of 107.6 μL. Each pool of samples was purified using 79 μL beads buffer (20% PEG-8000 in 2.5 M NaCL, final concentrations) and 50 μL Ampure XP Beads (Beckman Coulter, A63881), and eluted in 7 μL NF H2O.

Tagmentation was performed per pool by adding 5.5 μL double-stranded cDNA sample to 5.0 μL Nextera TD buffer (Illumina, 15027866), and 1.5 μL TDE1 Enzyme (Illumina, 15027865), incubated at 55°C for 5 min. Samples were directly placed on ice for at least 3 min. The reaction was terminated by adding 12 μL Buffer PB (QiaQuick, 19066) and incubating for 5 min at room temperature. Samples were purified using 48 μL Ampure XP beads and eluted in 10 μL NF H2O. Next, each sample was mixed with 2 μL P5 primer (10 μM), (5’-AATGATACGGCGACCACCGAGATCTACAC[i5]ACACTCTTTCCCTACACGACGCTCTTCCGATCT-3’; IDT), 2 μL P7 primer (10 μM) (5’-CAAGCAGAAGACGGCATACGAGAT[i7]GTCTCGTGGGCTCGG-3’; IDT), 6 μL NF H2O, and 20 μL NEBNext High-Fidelity 2X PCR Master Mix (NEB, M0541). Amplification was performed using the following program: 72°C for 5 min, 98°C for 30 sec, 15 cycles of (98°C for 10 sec, 66°C for 30 sec, 72°C for 1 min) and a final step at 72°C for 5 min. Samples were purified using 32 μL Ampure XP beads and eluted in 14 μL NF H2O. Libraries were visualized by electrophoresis on a 1% agarose and 1X TAE gel containing 0.3 μg/mL ethidium bromide (Invitrogen, 15585011). Gel extraction was performed to select products between 200 – 1000 bp using the Nucleospin Gel and PCR Clean-up kit (Macherey-Nagel, 740609). Samples were eluted in 15 μL NF H2O. cDNA concentrations were measured by Qubit dsDNA HS Assay kit (Invitrogen, Q32854). Product size distributions were visualized using Agilent’s Tapestation system (D5000 ScreenTape and Reagents, 5067-5588/9). Libraries were sequenced on the NextSeq 500 platform (Illumina) using a V2 75 cycle kit (Read 1: 18 cycles, Read 2: 52 cycles, Index 1: 10 cycles).

**Figure S1.**
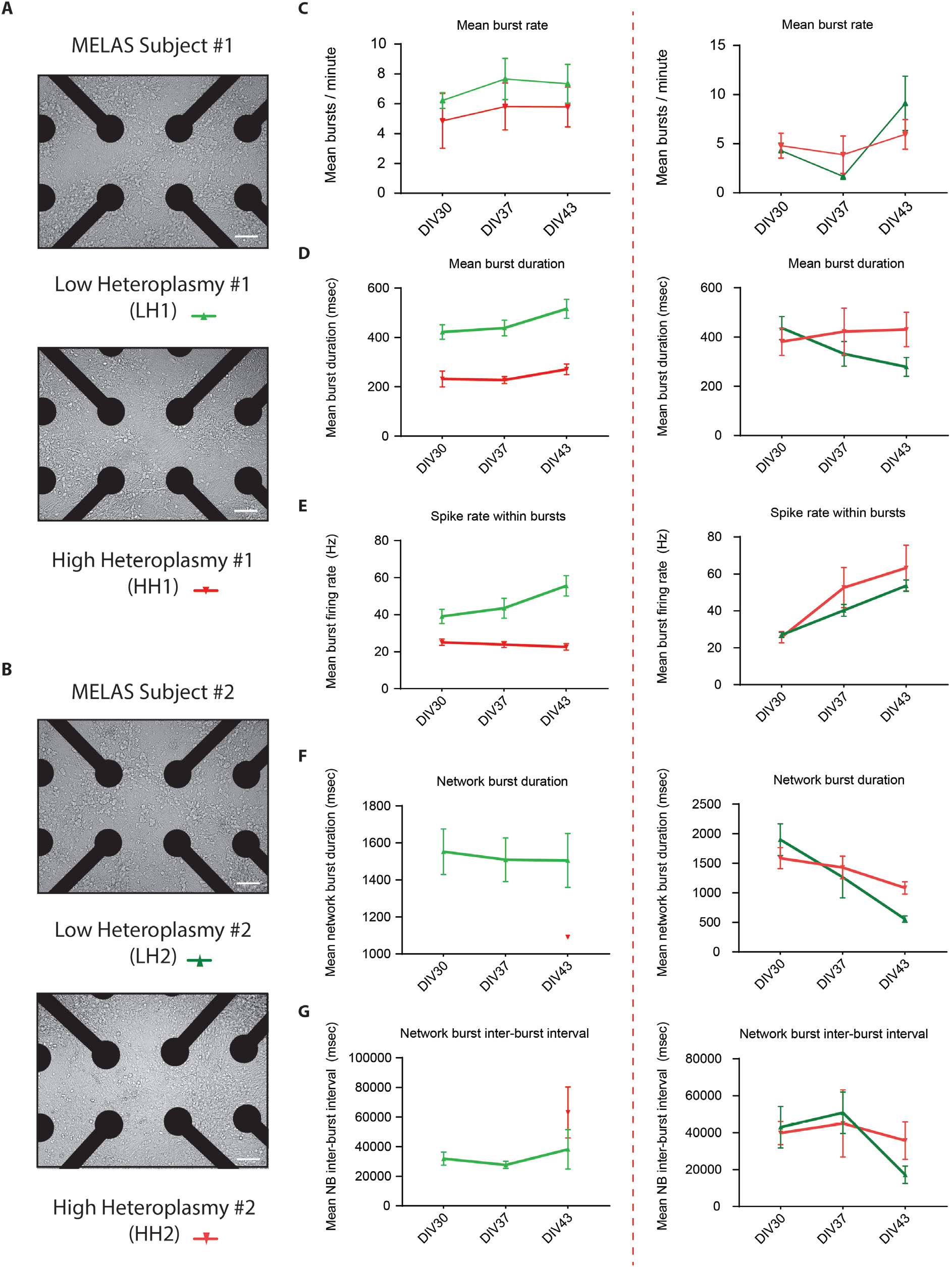
Density images and MEA data from the DIV30-43 developmental timeline, for lines LH1-2 and HH1-2. (A) Light fluorescence image showing density images for LH1 and HH1 (scale bar = 100 um), (B) Light fluorescence image showing density images for LH2 and HH2 (scale bar = 100 um). MEA networks were quantified based on (C) the mean number of bursts per minute (Mean burst rate), (D) the mean burst duration (msec), (E) the mean spike rate within bursts (Hz), (F) the mean network burst duration, and (G) the mean inter network burst interval (msec). Data represents means ± SEM. *P<0.05, **P<0.01, ***P<0.001, ****P<0.001, using restricted maximum likelihood model, with Holm-Sidak’s correction for multiple comparisons between treated and untreated samples.

**Figure S2.**
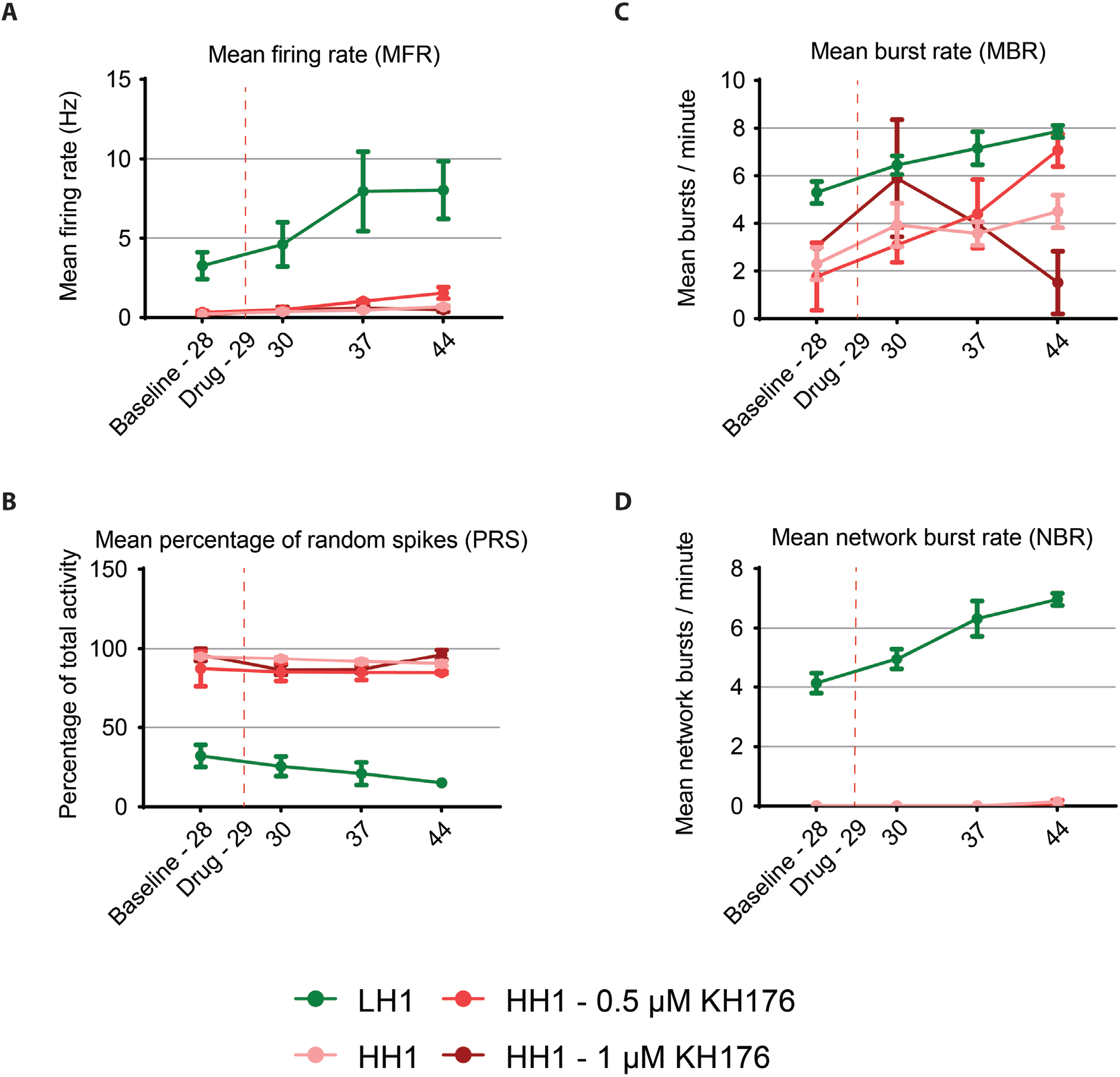
Principle component analysis on the neuronal- and astrocyte RNA samples. (A) Principal component analysis (PCA) plots displaying PC1 and PC2 for (A) the LH and HH neuronal samples, (B) the astrocyte samples co-cultured with LH or HH, (C) the LH1, HH1 and HH1+KH176 neuronal samples, and (D) the LH2, HH2 and HH2+KH176 neuronal samples.

**Figure S3.**
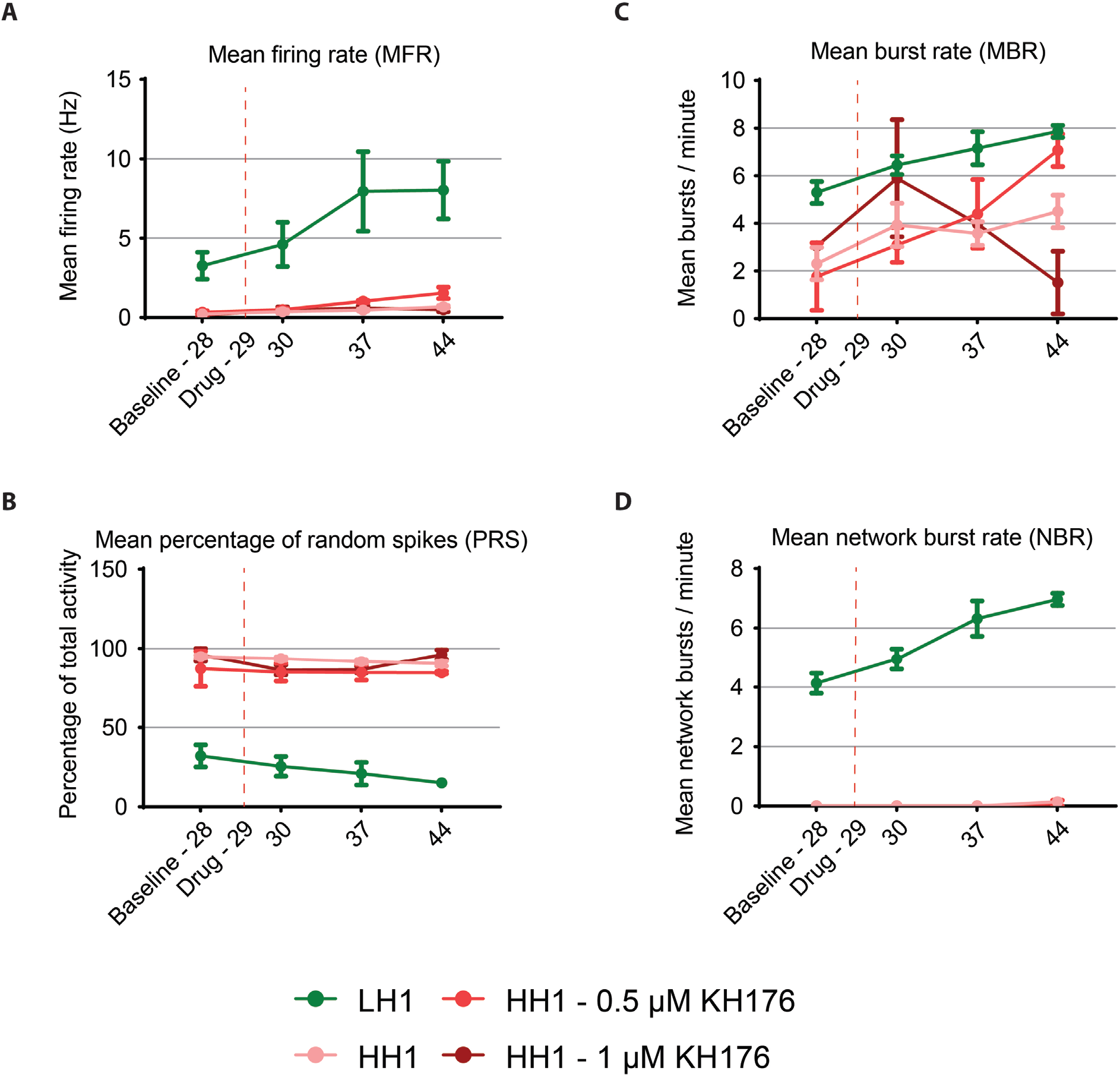
MEA data for sonlicromanol (KH176) treatment at mature network age (DIV29). HH1 cultures were treated from DIV29 onwards with either 500nm or 1000nm KH176 and recorded for 10 minutes of neuronal network activity on MEA; LH1 (n = 6), HH1 (n = 12), HH1 + 500nm KH176 (n = 4), HH1 + 1000nm KH176 (n = 4). We quantified (A) the mean firing rate (Hz), (B) the mean percentage of random activity (%), (C) the mean number of bursts per minute (burst rate), and (D) the mean number of network bursts per minute. Data represents means ± SEM. *P<0.05, **P<0.01, ***P<0.001, ****P<0.001, using restricted maximum likelihood model, with Holm-Sidak’s correction for multiple comparisons between treated and untreated samples.

**Figure S4.**
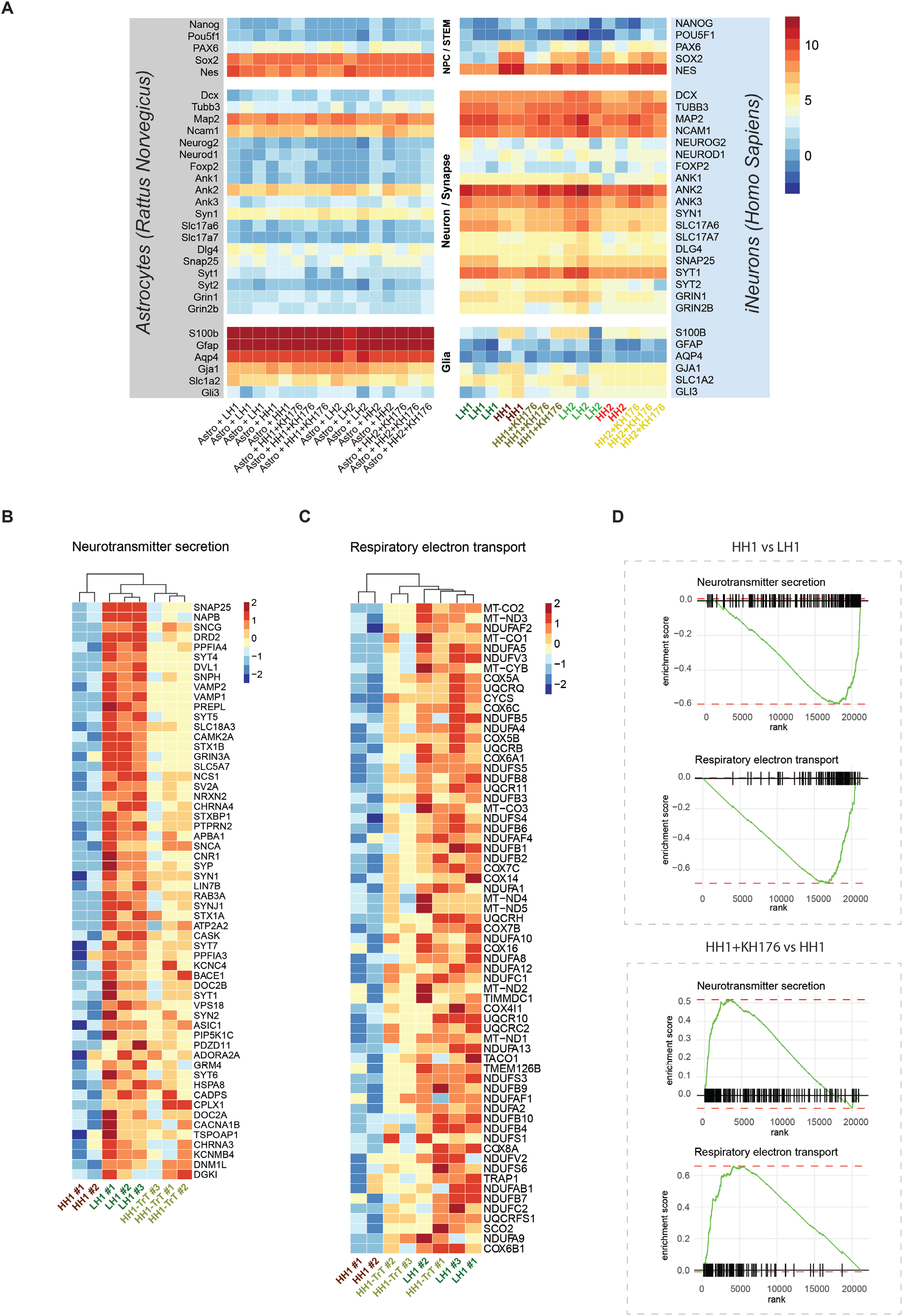
Heatmaps representing gene expression of full gene lists, and enrichment plots, for isogenic set LH1, HH1, and HH1+KH176. (A) Heatmap showing gene expression levels of NPC/stem cell genes (top section), neuronal- and synaptic genes (middle section), and glial genes (bottom section), including rodent homologs (left section) if expressed in astrocytes, and human homologs (right section) if expressed in iNeurons. Voom-transformed counts (log2 scale) corrected for batch effect and genetic background are shown. Heatmap showing overlapping leading edge genes (both for HH1 vs LH1, and for HH1+KH176 vs HH1) for (B) the neurotransmitter secretion gene set and (C) the respiratory electron transport gene set. Voom-transformed counts corrected for batch effect were scaled per gene. (D) Enrichment plots showing the running enrichment score for the neurotransmitter secretion gene set and the respiratory electron transport gene set, for comparisons of HH1 vs LH1 and HH1+KH176 vs HH1. Genes ranked on t-statistic from DE results are shown on the x-axis, with most up-regulated genes on the left and most down-regulated on the right.

**Figure S5.**
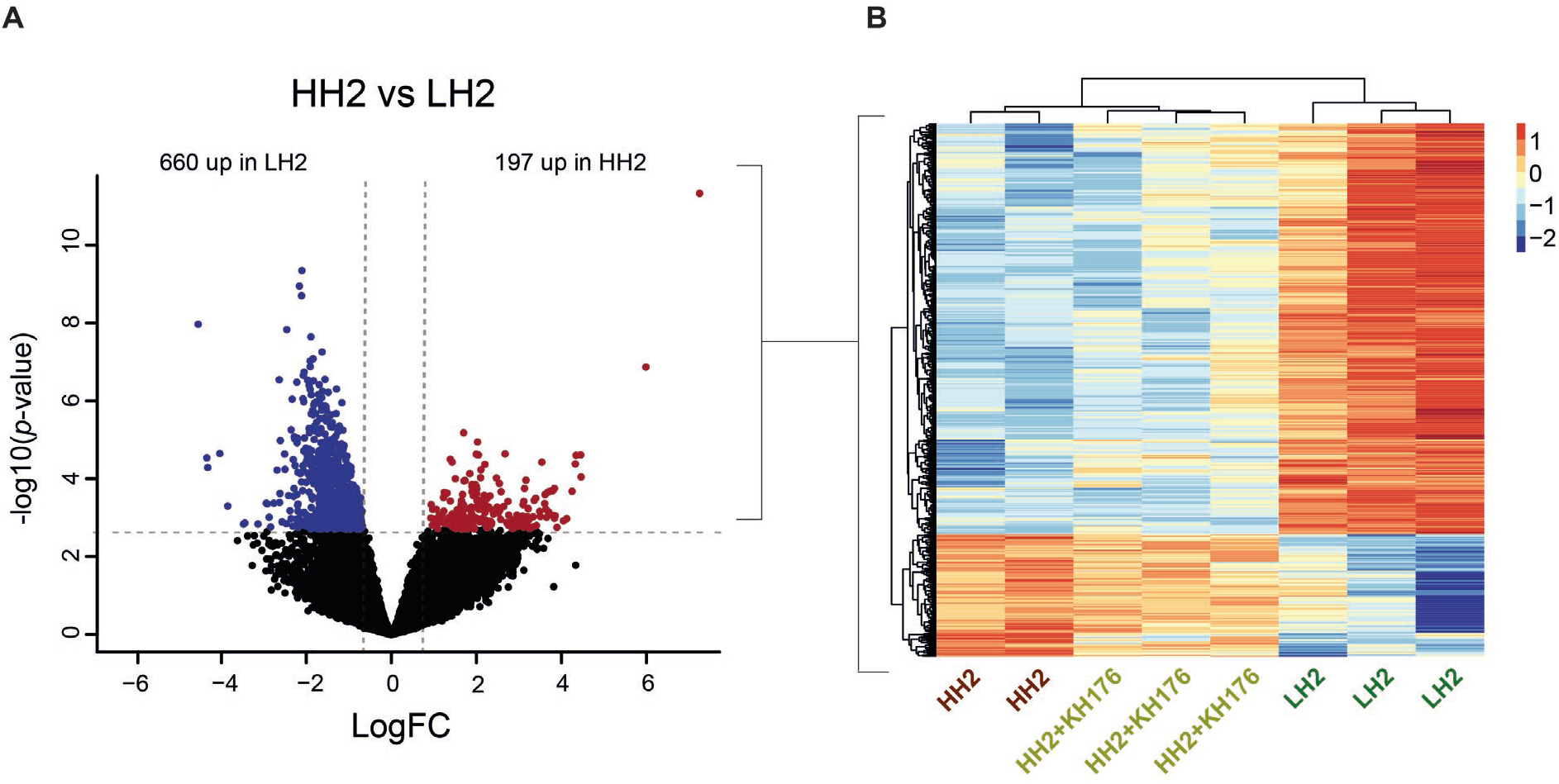
Comparing differential expression and GSEA for isogenic set #2, LH2, HH2, and HH2+KH176. (A) Volcano plot showing 857 differentially expressed (DE) genes (adj. *p* < 0.05) in HH2 compared to LH2 samples, with down-regulated genes in blue (log-fold change (LogFC) < 0) and up-regulated genes in red (LogFC > 0); 660 down-regulated and 197 up-regulated. (B) Heatmap showing expression of significant DE genes in HH2 vs LH2, for LH2, HH2 and HH2+KH176 samples. Voom-transformed counts corrected for batch effect were called per gene.

